# SUMO chains depolymerization induces slender to stumpy differentiation in *T. brucei* bloodstream parasites

**DOI:** 10.1101/2023.11.15.567218

**Authors:** Paula Ana Iribarren, Lucía Ayelén Di Marzio, María Agustina Berazategui, Andreu Saura, Lorena Coria, Juliana Cassataro, Federico Rojas, Miguel Navarro, Vanina Eder Alvarez

**Author notes:** Department of Life Sciences, Sir Alexander Fleming Building, Imperial College London, London, UK. Life Science Research Centre, Faculty of Science, University of Ostrava, 710 00 Ostrava, Czech Republic (Andreu Saura). These authors contributed equally to this work (Paula Ana Iribarren, Lucía Ayelén Di Marzio).

## Abstract

*Trypanosoma brucei* are extracellular protozoan parasites transmitted by tsetse flies that cause sleeping sickness in humans and nagana in cattle. Inside the mammalian host, differentiation from a bloodstream replicative slender form into a quiescent stumpy form allows the persistence of the parasite and the spread of the infection. SUMOylation is a reversible and dynamic post-translational modification of proteins that regulates diverse nuclear processes, such as DNA replication, repair and transcription. SUMO can be attached to its target proteins either as a single monomer or forming polymeric chains. We found that transgenic cell lines able to conjugate SUMO just as a monomer are attenuated *in vivo*. SUMO chain mutant monomorphic parasites display relapsing and remitting waves of parasitemia, at variance with wild-type parasites that cause unremitting parasitemia and mice death. Furthermore, when mice are infected with an analogous SUMO chain mutant generated in a differentiation-competent pleomorphic background, stumpy cells can be observed at unusually low parasitemia values. Our study reveals that SUMO depolymerization could represent a coordinated signal triggered during a quiescence activation program.

## INTRODUCTION

Pathogenic microorganisms have evolved multiple strategies to colonize, multiply and survive within their hosts. *Trypanosoma brucei* spp, is a protozoan parasite and the causative agent of sleeping sickness in humans and nagana in cattle [1]. This parasite lives extracellularly in the blood of the infected mammals, and one of its key mechanisms to succeed in generating long-term infections, is the antigenic variation of its major cell surface protein to evade the host immune adaptative response [2]. Another important feature is that the parasite can take control over its own proliferation, limiting the demand to the host, which otherwise would be lethal. This quorum sensing (QS) mechanism can induce the differentiation from a proliferative (slender forms, SL) to a quiescent stage (stumpy forms, ST) which is important not only to limit the parasitemia, but also to spread the infection, since differentiated cells are pre adapted for survival in a complete different metabolic niche, as it is the midgut of the tsetse fly vector [3].

The importance of a density-induced differentiation program can be largely appreciated when comparing the infection progress of laboratory adapted (monomorphic) *versus* naturally occurring (pleomorphic) parasites. Monomorphic parasites have been selected *in vitro* as actively dividing cells but are unable to control their population size *in vivo*. Thus, an infection with monomorphic parasites inexorably ends with the death of the animal. In contrast, pleomorphic parasites possess a density induced differentiation pathway, which allows the transition to an arrested form, and therefore can generate persistent infections.

Very recently, several components of this QS differentiation pathway have been identified [4], including the stumpy inducing factor (SIF) and its receptor [5, 6]. Downstream signalling might involve cascades through the activity of the kinases TOR and AMPK [7] eventually leading to G1/G0 arrest, mitochondrial activation, biochemical changes, and morphological transformation. However, full elucidation of the intracellular response is far to be completed. Phosphorylation, like many other reversible protein post-translational modifications (PTMs) enables fast modulation of protein function in response to environmental conditions. In this work, we have unveiled a connection between another PTM, i.e. SUMOylation, and the differentiation process that takes place inside the mammalian host.

SUMOylation is a PTM conserved in eukaryotic organisms involving the covalent attachment of the Small Ubiquitin-like MOdifier (SUMO) to internal lysine residues within target proteins [8]. This modifier usually alters the interaction surface of its substrates, controlling the biological activity, stability, or subcellular localization, among other possible outputs. SUMOylation is essential in *T. brucei* [9, 10] and regulates many important biological processes, as inferred by the proteomic analysis of its target proteins [11]. Interestingly, in *T. brucei* bloodstream forms (BSF) SUMO is enriched in a particular region of the nucleus, colocalizing with the RNA polymerase I at the expression-site body (ESB) within the active variant surface glycoprotein (VSG) expression site (VSG-ES). This highly SUMOylated focus (HSF) creates a permissive environment for VSG transcription [12].

Like ubiquitin, SUMO can covalently modify substrates as a monomer or by forming different types of chains through internal lysine residues within the N-terminal of SUMO, conferring a new level of versatility and complexity to this dynamic PTM. SUMO chains have been described in many eukaryotic organisms, participating for example in mitosis [13] or regulating the formation of promyelocytic leukemia nuclear bodies [14] and the synaptonemal complex [15]. We have demonstrated that *Tb*SUMO is able to form polymeric chains through its K27 and that these structures are important for the assembly of nuclear foci, suggesting a regulatory role of polySUMOylation in chromatin organization in procyclic forms (PF) [16, 17].

In this work, we have examined the role of SUMO chains in BSF using transgenic monomorphic cell lines unable to assemble them. We found that, at variance with the virulent monomorphic parental strain, SUMO chain mutants can establish persistent infections in mice. This behavior is related with a “stumpy-like” characteristic of the parasites since mutants displayed some typical stumpy markers and an increased differentiation kinetic from BSF to PF induced by *cis*-aconitate. Thus, in contrast with the “blind to the signal” status of the parental monomorphic cell line, the absence of SUMO chains renders the BSF sensitive to the population size. To confirm our results, we generated a similar mutant but using pleomorphic cells able to differentiate naturally. We found that *in vivo* the parasites become arrested at lower cell densities exhibiting a typical stumpy morphology and expressing the specific marker PAD1. We propose that SUMO chain dynamics influence the ability to differentiate from SL to ST forms, being polySUMOylation a SL retainer signal that can be relieved after its debranching.

## RESULTS

### SUMO chain mutant monomorphic parasites growth normally *in vitro* but have a reduced virulence in mice

To investigate the role of SUMO chains in monomorphic BSF of *T. brucei,* we generated a mutant cell line in which all internal lysine residues in SUMO were replaced with arginine (*Tb*SUMO*all*KR, Figure 1A). Since SUMO is essential in *T. brucei*, we first evaluated whether the absence of chains could have a detrimental effect. SUMO*all*KR can be processed and conjugated to target proteins showing a reduced intensity in the SUMOylation pattern by western blot (Figure 1B) but without presenting any growth phenotype in culture (Figure 1C). Chromatin SUMOylation at the active VSG expression site is a major feature of BSF [12]. The typical SUMO labeling analyzed by indirect immunofluorescence microscopy (IF) consists of a diffuse nuclear pattern and one highly SUMOylated focus (HSF) in ∼80% of the cells. In contrast, in the SUMO chain mutant parasites, the signal was detected both in the nucleus and the cytoplasm with a more dispersed distribution (less than 40% of the cells presenting a HSF) (Figure 1D). This difference, however, does not impact on VSG mRNA or protein levels (Figure S1), suggesting that mono and/or multiSUMOylation are sufficient to promote VSG expression.

**Figure 1:**
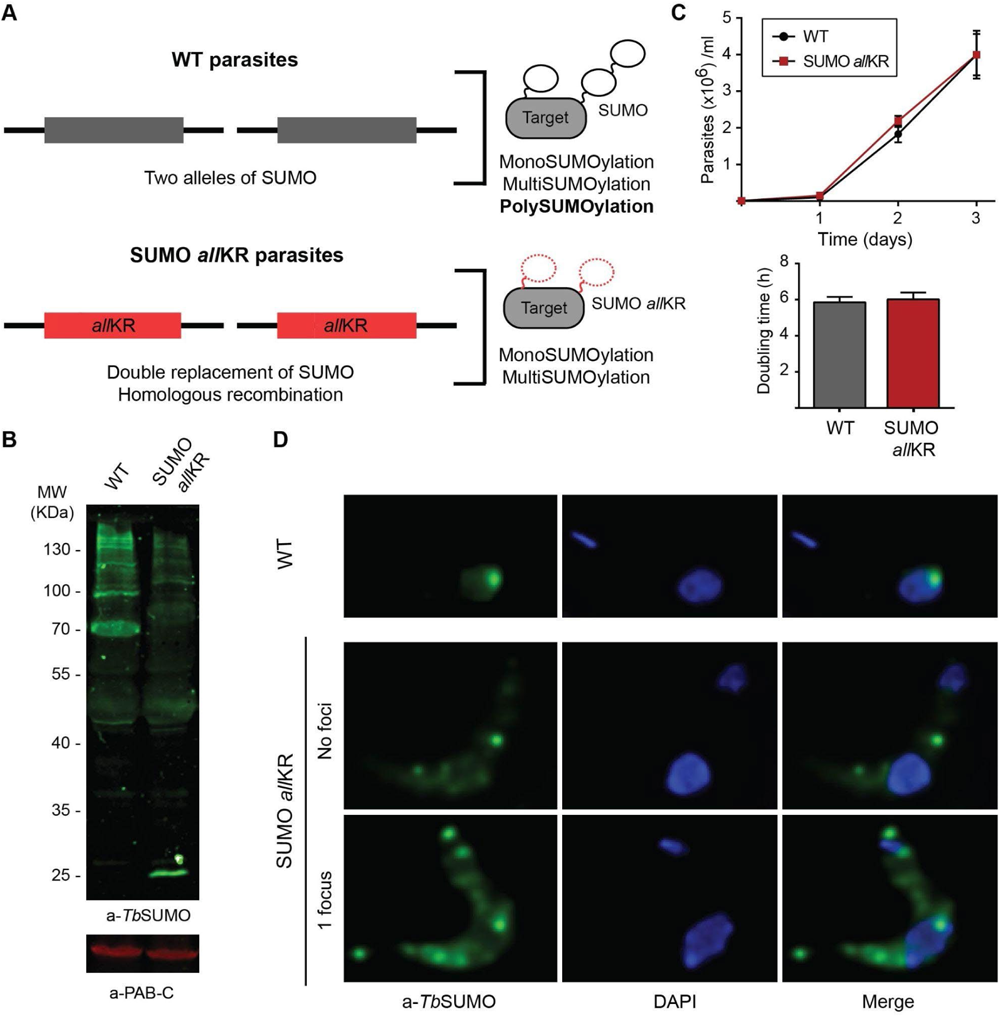
Generation of SUMO chain mutant monomorphic BSF parasites. **(A)** Schematic representation of the generation of *Tb*SUMO *all*KR parasites. **(B)** Growth curves for SUMO *all*KR and wild type (WT) parasites. WT and transgenic parasites were cultured up to one month without observing significant differences in growth rate. Doubling time was calculated by daily subculture back to 1 × 10^5^/ml to maintain log-phase growth. **(C)** Conjugating ability of WT *Tb*SUMO (WT) and SUMO *all*KR parasites. Parasites were boiled in Laemmli’s sample buffer immediately after harvesting. Proteins were separated by electrophoresis using a 10% SDS-poliacrylamide gel (3×10^7^ cells/lane). SUMO conjugates were analyzed by Western blot using anti-*Tb*SUMO antibodies and anti-PAB-C antibodies as loading control. **(D)** Double indirect three-dimensional immunofluorescence (3D-IF) analysis of WT and SUMO *all*KR BF parasites. Nuclear and kinetoplast DNA were visualized by DAPI staining (blue). Representative images of anti-*Tb*SUMO (green) and anti-*Tb*SUMO-DAPI merged images are shown.

We next investigated the influence of SUMO chains *in vivo* using a mouse model of infection (Figure 2). Inoculation of wild-type (WT) parasites was lethal to 100% of mice within 6 days, as expected for a monomorphic cell line. In contrast, infection with SUMO chain mutant parasites resulted in a markedly prolonged survival: 86% at day 6-post infection (dpi) and 43% at 10 dpi. This phenotype was reverted when the mutant was complemented with a WT copy of SUMO, killing all mice at 7 dpi, like animals infected with SUMO chain competent parasites (Figure 2A). To further study this, we analyzed parasitemia daily. For WT monomorphic BSF, parasitemia was first detected by 4 dpi and increased approximately 100-fold per day thereafter (n=6). All mice developed a single wave of unremitting parasitemia with a terminal outcome at day 6 (Figure 2B, panel WT and Figure S2). In contrast, mice infected with SUMO chain mutants (n=17) exhibited two or three waves of parasitemia and in some cases even cleared the infection (Figure 2B, panel *all*KR and Figure S2).

**Figure 2:**
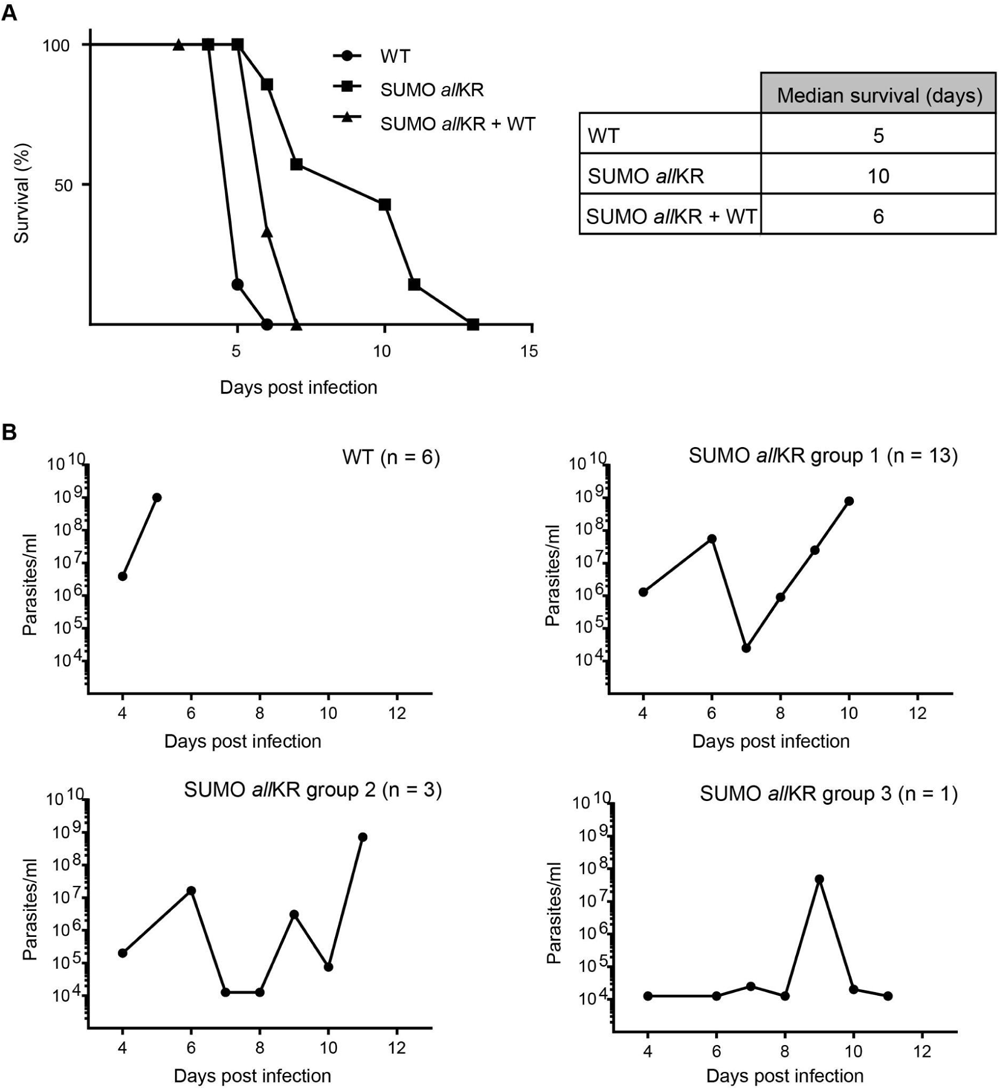
Mice infections with SUMO chain mutant monomorphic BSF parasites. **(A)** BALB/c mice were infected intraperitoneally with 5000 wild type (WT; circles), SUMO *all*KR (squares) or chain mutant parasites complemented with a WT copy of SUMO (SUMO *all*KR+WT; triangles). Survival was monitored daily and is shown by a Kaplan-Meier curve. **(B)** Time course of parasitemia in mice infected with WT or SUMO *all*KR parasites. Animals were grouped according to their different parasitemia profile.

### Monomorphic SUMO chain mutant parasites display stumpy-like characteristics

After ruling out potential differences in the humoral immune response of the host (Figure S3) as well as in the ability of the parasites to internalize and degrade anti-VSG antibodies (Figure S4), we hypothesized that the absence of polySUMOylation could be restoring, at least partially, the original ability of the SL form to differentiate to the ST. At the peak of the parasitemia, we observed cells that cannot be recognized as genuine stumpy parasites based on their morphology or biochemical markers (not shown) [18]; however, we found a higher proportion of 1K 1N cells (Figure 3A) and increased mRNA levels for the characteristic stumpy markers PAD1 and PAD2 when compared to WT cells (Figure 3B).

**Figure 3:**
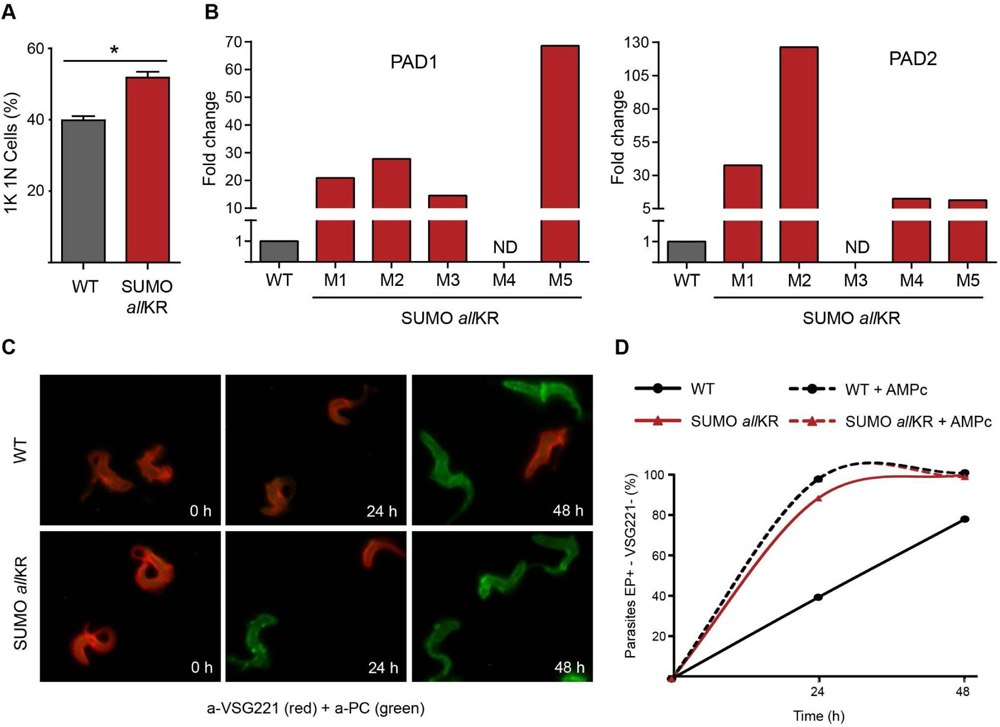
Analysis of stumpy markers in monomorphic SUMO chain mutants. **(A)** Blood smears from the first peak of parasitemia were stained with DAPI for nucleus and kinetoplast configuration analysis (**, p<0.01). **(B)** Quantification of RNA transcript levels of the characteristic stumpy markers PAD1 and PAD2 was performed by qRT-PCR from RNA samples derived from mice infections. WT and SUMO *all*KR parasites were isolated at the first peak of the parasitemia. Samples were normalized against 7SL. **(C)** *In vitro* bloodstream-to-procyclic differentiation induced by cis-aconitate (CA) was analyzed in SUMO *all*KR and wild type (WT) parasites. Differentiation to the procyclic form was triggered in SDM-79 medium at 28°C using 6 mM CA. The differentiation process was evaluated following changes in the expression of stage-specific surface markers (VSG221 and procyclin) for 48 h by indirect immunofluorescence. Representative images of anti-VSG221 (red)-anti-EP (green) merged images are shown. One out of five representative experiments is shown. **(D)** WT + 8-pCPT-cAMP and SUMO *all*KR + 8-pCPT-cAMP parasites were pre-treated with the cAMP analogue 8-pCPT-cAMP [8-(4-Chlorophenylthio) adenosine 3′,5′-cyclic monophosphate] during 24 h before shifting the temperature and adding CA.

To investigate the presence of stumpy-like forms, we challenged the parasites with high concentrations of *cis*-aconitate (CA) at low temperature to stimulate the differentiation to PF, which is thought to occur through this intermediate stage [19, 20]. During this transformation, the VSG coat is released and replaced by a different coat consisting of invariant GPEET and EP procyclins. For pleomorphic cell lines containing short stumpy cells, this process is synchronous, and it only takes 12 h to complete. In contrast, for monomorphic cells, the process is non-synchronous and takes 36-48 h to be completed. To assess the ability of monomorphic, SUMO chain competent or SUMO chain mutant parasites, to differentiate into PF, we monitored the switching of the coat by IF analysis (Figure 3C). As expected, transformation of SUMO chain competent cells occurred with slow kinetics. After 24 h of CA treatment about 60% of the cells were still expressing VSG and the number decreased to 28% 24 h later. In contrast, SUMO chain mutants displayed considerably higher rates of differentiation. Within 24 h, ∼90% of the cells were already EP positive and VSG negative, and the transformation was almost complete at 48 h. Interestingly, these differences were abolished when the parasites were pre-treated with the cAMP analogue 8-pCPT-cAMP [8-(4-Chlorophenylthio) adenosine 3′, 5′-cyclic monophosphate] (a compound able to induce slender to stumpy differentiation [21]) during 24 h before shift temperature and CA addition (Figure 3D).

Altogether these results suggest that the absence of polySUMOylation could stimulate the generation of stumpy-like cells rendering monomorphic bloodstream parasites more sensitive to CA-triggered differentiation and that the underlying mechanism is upstream of AMP signalling.

### SUMO chain mutant pleomorphic parasites are primed for stumpy differentiation

To properly address the role of SUMO chains depolymerization in ST formation, we generated a similar SUMO chain mutant but in differentiation-competent pleomorphic parasites. We first analyzed the SUMOylation pattern by western blot and, consistent with an impairment of chain formation, a significant reduction in the high molecular weight adducts was observed (Figure 4A). When the growth profile was assayed (Figure 4B), a small effect on the duplication time was found. This growth phenotype is linked to an accumulation of non-dividing cells as shown by the increase in the number of cells with 1K 1N configuration (Figure 4C). To analyze the generation of ST *in vitro*, we exposed the parasites to an increased local concentration of SIF by culturing them on HMI-9-agarose plates for 4 days followed by immunodetection of the stumpy marker protein PAD1. As shown in Figure 4D and 4E, chain mutant parasites showed a significantly higher level of cells expressing PAD1 (74% ± 2%), whereas only 35% ± 6% of the WT cells exhibited expression of PAD1 at the same time-point. Furthermore, while PAD1 was already observed in the surface of the chain mutant parasites, in WT cells it was observed in vesicles and the flagellar pocket, very likely trafficking to the membrane. Finally, chain mutants exhibited an increased rate of transformation to procyclic forms when exposed to the differentiation signal *cis*-aconitate (Figure S5).

**Figure 4:**
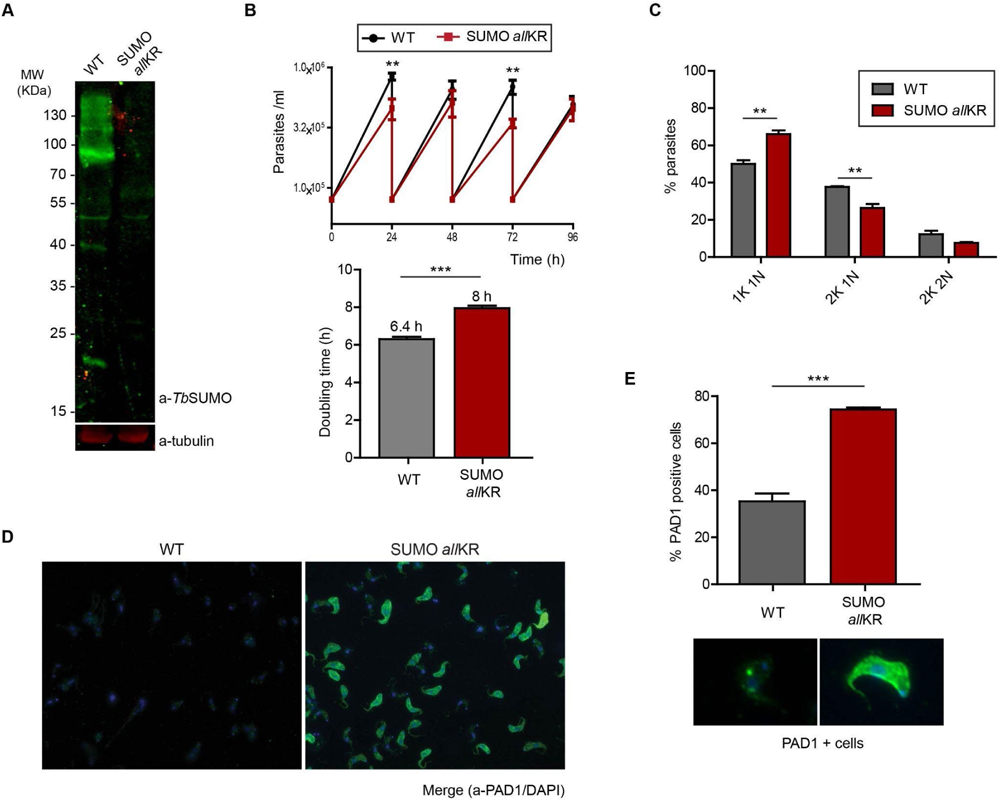
Generation of SUMO chain mutant pleomorphic parasites. **(A)** SUMOylation pattern in pleomorphic parasites. Whole-cell extracts of WT or SUMO *all*KR parasites were boiled in Laemmli sample buffer immediately after harvesting. Proteins were separated by SDS-PAGE and transferred to a nitrocellulose membrane, followed by immunoblotting with anti-SUMO antibodies. Tubulin was used as loading control. **(B)** Growth curves of pleomorphic parasites. WT or SUMO *all*KR parasites were cultured in vitro under regular conditions in HMI-9. Doubling time was calculated by daily subculture back to 1×10^5^/ml to maintain log-phase growth (**, p<0.01; ***, p<0.001). **(C)** Nucleus and kinetoplast configurations. Pleomorphic parasites in exponential growth phase were fixed and stained with DAPI for nucleus and kinetoplast configuration analysis (**, p<0.01). **(D)** Stumpy formation in HMI-9 agarose plates. WT or SUMO *all*KR parasites were plated onto semi solid agarose plates. After 4 days, parasites were harvested, fixed and stained with DAPI (blue) and anti-PAD1 antibodies (green). **(E)** Quantification of parasites from (D) (***, p<0.001). Representative images of PAD1 positive cells are shown.

Having confirmed that the absence of SUMO chains renders the parasite more susceptible to the QS signal *in vitro*, we monitored the consequences in mice infections. Figure 5A and Figure S7 demonstrates that infection with SUMO*all*KR pleomorphic parasites resulted in consistently reduced parasitemia and prolonged host survival (Figure 5B). This infection pattern is driven by a premature growth arrest (Figure 5C) since >95% of SUMO *all*KR parasites were 1K1N configuration at the first peak when the parasitemia was as low as 10^6^ parasites/ml. Even at these low densities, we observed that parasites were morphologically stumpy and displayed the stumpy-specific marker protein PAD1 on their surface (Figure 5D and 5E) at variance with the wild-type pleomorphic cells. Thus, our results clearly demonstrate that the absence of SUMO chains is a signal for stumpy development during infections.

**Figure 5:**
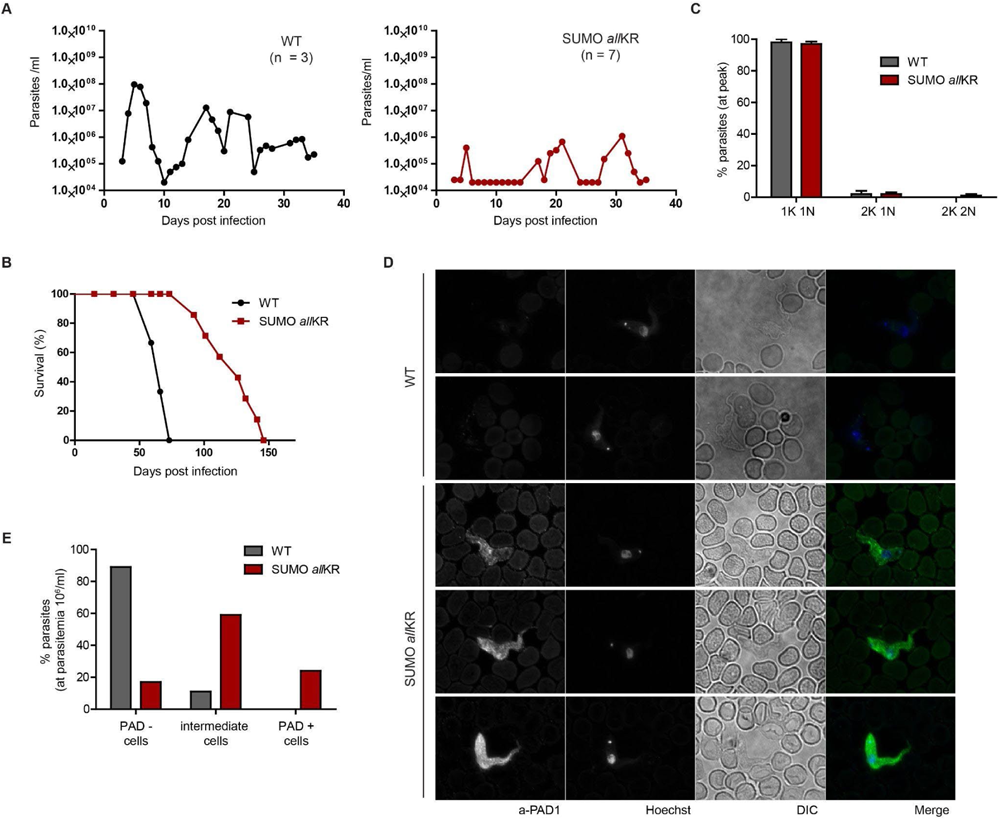
Mice infections with SUMO chain mutant pleomorphic parasites. Mice infections with pleomorphic parasites. BALB/c mice were infected intraperitoneally with 5000 WT or SUMO *all*KR parasites. **(A)** Time course of parasitemia in mice infected with WT or SUMO *all*KR parasites. **(B)** Mice survival was monitored daily and is shown by a Kaplan-Meier curve. **(C)** Nucleus and kinetoplast configurations. Methanol-fixed blood smears of infected mice on the peak of parasitemia were prepared and stained with Hoechst. **(D)** Methanol-fixed blood smears of infected mice were prepared and stained with anti PAD1 antibodies (green) and Hoechst (blue). **(E)** Quantification of PAD1 stained parasites of (D).

### SUMO chain dynamics in pleomorphic parasites

Next, we looked if any changes in the mRNA abundance of the main SUMO deconjugating enzyme *Tb*SENP [16] have been reported for the differentiation competent AnTat1.1 cells during synchronous transformation [22]. Consistent with our hypothesis, the mRNA profile showed that the highest levels of SENP were reached at the stumpy forms derived from infections in mice ([22], Additional file 3b).

Then, we investigated if this increase in *Tb*SENP expression during stumpy formation could correlate with any difference in the labelling of SUMO. For this, we induced differentiation with 8-pCPT-cAMP and assessed transformation using PAD1 antibody as a stumpy marker. For PAD1 negative cells, we observed the typical nuclear foci of SUMOylated proteins for the majority of the cells (Figure 6A, 87.5% in foci, 12.5% dispersed). In contrast, PAD1-positive cells displayed a predominantly diffuse labelling for SUMO (58% of the cells with dispersed signal versus 42% of the cells with visible foci), resembling the one described for the monomorphic chain mutant cell line (Figure 6B). Taken together, these results further support the notion that debranching SUMO is indeed priming bloodstream parasites for subsequent differentiation.

**Figure 6:**
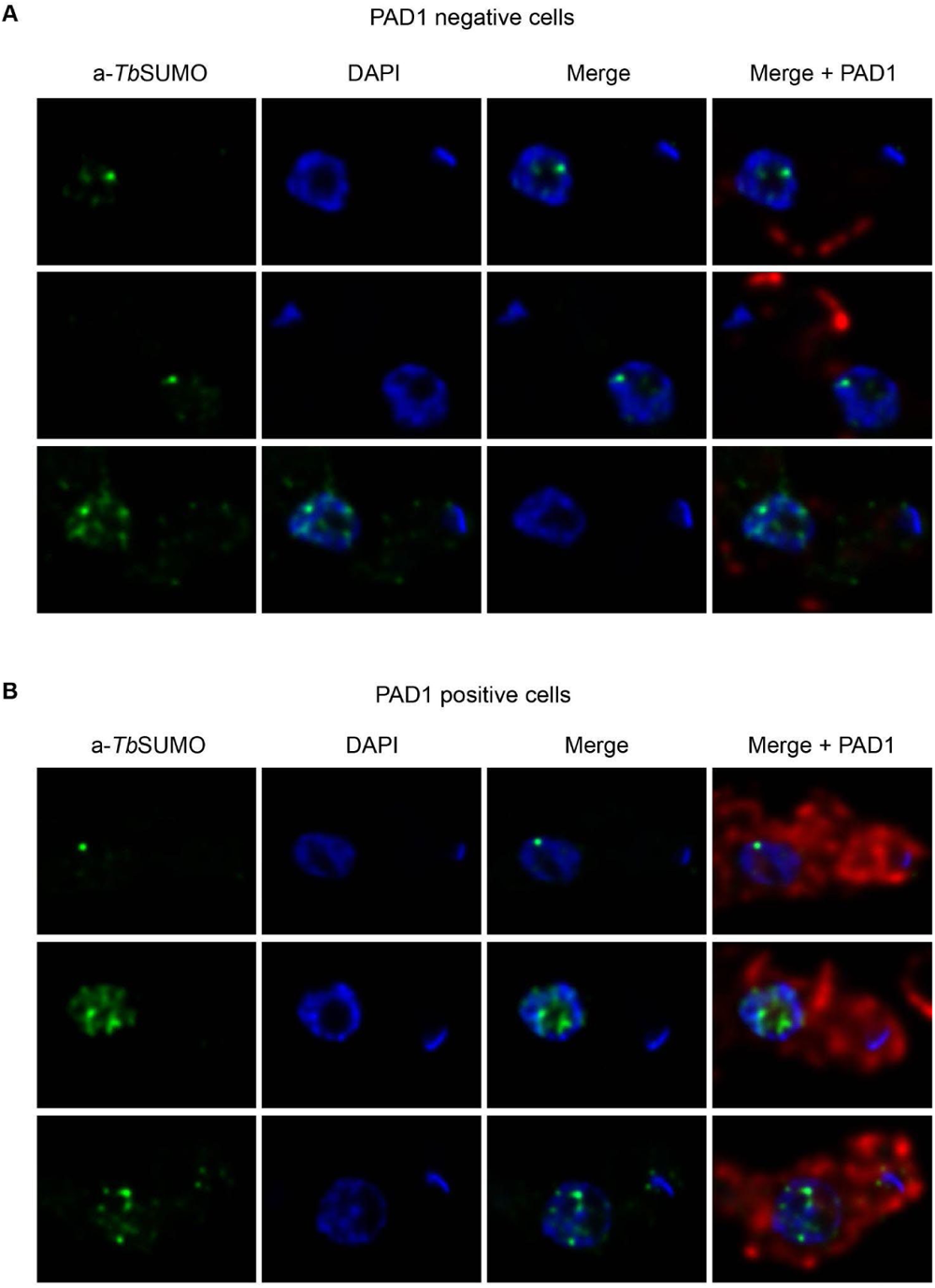
SUMO chain dynamics in pleomorphic parasites. Double indirect three-dimensional immunofluorescence (3D-IF) analysis of pleomorphic AnTat 90:13 BF cultured at low density (2 x 10^5^ cells/ml) and incubated with 1μM 8-pCPT-cAMP during 18 h. Nuclear and kinetoplast DNA were visualized by DAPI staining (blue). Representative images of anti-*Tb*SUMO (green), anti-PAD1 (red) and merged images are shown. **(A)** PAD1 negative cells. **(B)** PAD1 positive cells.

## DISCUSSION

The results obtained in this work reveal that SUMO chains in *T. brucei* BSF constitute an inhibitory signal for the differentiation from slender to stumpy, and that the disassembly of these chains in a context of an infection can promote the persistence of the parasite in the host. Our conclusion is based on several experimental findings: 1) the outcome of the infection with SUMO chain mutants BSF; 2) the expression of stumpy markers when parasites reach the peak of the parasitemia; 3) the accelerated rate of differentiation to PF after *cis-*aconitate exposure and 4) the significant difference in the SUMO pattern observed by IF in pleomorphic parasites.

In this study, we imitated the monoSUMOylation status of target lysine residues, abolishing the ability of SUMO to polymerize. For this purpose, both endogenous SUMO alleles were replaced by a version in which all lysine residues are mutated to arginine (SUMO *all*KR), thus preserving the net charge but hindering further internal modification of SUMO with SUMO. We have chosen this strategy over specific mutation of the lysine residue at position 27, as previous reports had shown that in the absence of the major acceptor lysine residue some other lysines (otherwise cryptics) might be able to undergo modification, in some cases with the assistance of SUMO ligases [23–29]. This phenomenon has been also observed for ubiquitin [30, 31]. Considering this and taking into account that we wanted to analyze the effect of the chain deficiency, we decided to replace all lysine residues to be sure that chains were absent.

SUMO chains are not essential for BF axenic growth, at least under our experimental conditions using routine HMI-9 culture medium. Of note, the SUMO chain mutant parasites express average levels of VSG, suggesting that monoSUMOylation of certain factors is sufficient for PolI recruitment to the active expression site. The most surprising phenotype for either monomorphic or pleomorphic mutant cell lines was detected *in vivo*. We consistently observed that the abrogation of SUMO chains in *all*KR parasites resulted in reduced virulence and increased survival of the host.

For monomorphic parasites, mice infected with SUMO chain mutants presented relapsing and remitting waves of parasitemia, reaching densities not higher than 10^8^/ml and clearing the day after. This behaviour is completely unusual for a monomorphic cell line since these developmentally-incompetent parasites cannot control parasitemia through differentiation to quiescent stumpy cells, killing the host within the first days post infection as can be observed for the parental WT parasites. Pleomorphic strains, on the other hand, are able to complete the parasite natural life cycle showing persistent infections in animal models, with undulating parasitemia [3, 32, 33]. Thus, it seems that the abrogation of SUMO chains has reverted the typical behaviour of monomorphic strains *in vivo*, acquiring characteristics of pleomorphic parasites, with prolonged mice survival and attenuated parasitemia. To our knowledge, this characteristic growth pattern has been reported for only one other monomorphic strain which was a null mutant for the calflagin gene [34], although the underlying mechanism was not further investigated.

After discarding potential differences in the humoral immune response of the host as well as in the ability of the parasites to internalize and degrade anti-VSG antibodies, we hypothesized that the altered course of the infection could be the result of the differentiation to stumpy-like cells. Indeed, the expression of the characteristic stumpy markers PAD1 and PAD2 were increased at the mRNA level (but not at the protein level) when compared to the control WT monomorphic parasites. These intermediate phenotypes, that have not fully completed the developmental process, have been previously described, expressing PAD1 mRNA prior to protein expression and final morphological transformation to mature stumpy cells [33, 35–40]. Notably, these intermediate forms were observed only *in vivo*, while *in vitro*-cultured SUMO *all*KR parasites probed to be negative for all of the assays used to evaluate stumpy characteristics (morphology, mild acid resistance, mitochondrial activity, cell cycle status, expression of stumpy markers; data not shown) [18] except for differentiation using CA stimulus. In this latter case, SUMO chain mutants displayed an accelerated kinetics of differentiation to PF compared to control parasites, showing a significantly increased replacement of the VSG coat by procyclin 24 h post induction. Furthermore, this difference was abolished by treatment with a cAMP analogue, suggesting that SUMO chain signalling might be upstream of the AMPK pathway [7]. Taken together these results suggest that, unlike WT monomorphic strains, the absence of polySUMOylation renders the parasites more susceptible to differentiation while a host-parasite interaction is required to trigger stumpy-like cells. In agreement with this, Rojas et al. have recently described the importance of the interplay between the environmental information and trypanosomes being both, host and parasite proteins, required for quorum sensing signaling in developmentally competent cells [5].

We confirmed our hypothesis generating a similar mutant in a differentiation competent pleomorphic background. Mutant parasites grown on HMI-9 agarose plates displayed all typical stumpy form features, including PAD1 protein expression at the cell surface, arrest at the cell cycle G_0_-G_1_ phase and prompt differentiation to procyclic forms when incubated with *cis*-aconitate. Furthermore, *in vivo,* these mutants differentiated into stumpy forms at low densities (around 1 10^6^ - 1 10^7^ parasites/ml) with a concomitant reduced parasitemia and prolonged survival of the infected mice.

When analyzing the subcellular localization of SUMO in pleomorphic parasites, a dispersed distribution was observed in a large number of cells. This distribution is in deep contrast to the typical distribution for SUMO in WT monomorphic parasites, with one or two HSF in most cells [12] but resembles the pattern observed for the SUMO chain mutant monomorphic cell line. Then, it is possible to speculate that SUMO debranching and differentiation might be coordinated events, being polySUMOylation involved in this process as a negative regulator during natural infections. In fact, the differentiation of pleomorphic parasites is accompanied by a significant increase in the mRNA of the tightly regulated and main deSUMOylase [16]. In this context, it would be interesting to uncover the identity of the substrates responsible for this phenotype.

SUMO chains have been implicated in several processes, in particular, it has been noticed that they play a crucial role when cells are under stress [41]. The best-characterized example is during proteotoxic stress, where they act as a protein degradation signal targeting polySUMOylated proteins for proteolytic ubiquitination [42]. In addition, other studies [43, 44] have proved that the SUMO system is involved in transcriptional control, and one particular study showed that SUMO chains are responsible for the activation of survival pathways during stress through transcriptional derepression of stress-regulated genes [45]. Up to date, site-specific-proteomic studies have allowed the identification of SUMO targets in PF and BF parasites, many of them associated with relevant nuclear processes, such as DNA replication and repair, RNA metabolism, transcription and chromatin remodelling, among others [11] (Saura et al, *under review*). Considering that during slender to stumpy transition there is a downregulation in transcription and protein synthesis [18], some reported SUMO-modified chromatin remodelers might constitute interesting candidates to address their polySUMOylation status regarding differentiation.

Based on our results, we propose that polySUMOylation of specific substrates is important to retain *T. brucei* in a slender form while debranching of SUMO leads to stumpy differentiation following quorum sensing signalling upon host-parasite interaction. We believe that SUMO chain dynamics constitute an additional level of control of this PTM and a novel layer that can be modulated to influence parasite differentiation.

## MATERIALS AND METHODS

### Trypanosome culture

*Trypanosoma brucei* bloodstream form (BF) “Single Marker” (SM) parasites (T7RNAP TETR NEO), pleomorphic BF EATRO 1125 [46] and transfected cells were grown at 37°C and 5% CO2 in HMI-9 media [47] (Life Technologies, Carlsbad, CA, USA) supplemented with 10% (vol/vol) heat-inactivated fetal calf serum (Natocor, Córdoba, Argentina) and appropriate antibiotics.

### Generation of SUMO chain mutant parasites

For SUMO *all*KR strain, we employed a synthetic construct (GenScript, Piscataway, NJ, USA) described previously [17]. This construction contained the coding sequence for 8 histidine residues, an hemagglutinin (HA) tag, the complete open reading frame (ORF) of *Tb*SUMO (Tb927.5.3210) with all its 8 lysine residues replaced by arginine residues, a 200 bp fragment of the 5’end of 3’untranslated region (UTR) of the gene, a *Xho*I restriction site and 250 bp of the 3’end of *Tb*SUMO 5’UTR. This was cloned into the endogenous locus tagging vector pEnT6 [48], with puromycin or hygromycin resistance marker cassette, to allow the sequential replacement of *Tb*SUMO alleles by homologous recombination through its UTRs. The same vector, but with WT *Tb*SUMO ORF [11] and blaticidin resistance marker cassette, was employed to reverse SUMO *all*KR phenotype. Vectors were linearized with *Xho*I (New England Biolabs, Ipswich, MA) and used to transfect BF parasites based on the protocol described by the Cross laboratory (http://tryps.rockefeller.edu/). For the SUMO *all*KR pleomorphic strain, one allele of *Tb*SUMO was first replaced by a blasticidin resistance marker cassette. To achieve this, a pGEM-T Easy vector (Promega, Madison, WI, USA) with the blasticidin cassette flanked by a fragment of the 5’ UTR and 3’ UTR of the SUMO gene, to allow replacement by homologous recombination, was made. This vector was linearized with *NotI* (New England Biolabs, Ipswich, MA) and used to transfect BF parasites. After confirming the correct replacement of the blasticidin cassette, this SUMO hemi KO line was transfected with the *Tb*SUMO *all*KR construct, as described for the monomorphic strain. Log phase cells (2-3 ×10^6^ ml^−1^) collected by centrifugation were resuspended in 90 µl of Tb-BSF buffer (90 mM Na_2_HPO_4_, 5 mM KCl, 0.15 mM CaCl_2_, 50 mM HEPES, pH 7.3) and mixed with 10 µl of linearized DNA (5-15 µg) in a 0.2 cm electroporation cuvette (BTX, Harvard Apparatus, Holliston, MA, USA). Parasites were then subjected to one pulse using X-001 program in the Amaxa Nucleofector 2b (Lonza Cologne AG, Germany). Transfected cells were cloned after 6 h in 24-well dishes with the appropriate selective drugs (2.5 μg/ml of G-418; 0.1 μg/ml of puromycin; 5 μg/ml of hygromycin; 5 μg/ml of blasticidin (InvivoGen, San Diego, CA, USA)). The correct replacement of WT *Tb*SUMO alleles in SUMO *all*KR clones was confirmed by Polymerase Chain Reaction (PCR) using specific primers, as described previously [17]. *Tb*SUMO PCR products were sequenced (Macrogen, Seoul, Korea).

### Electrophoresis and immunoblotting

Parasites collected by centrifugation were resuspended in Laemmli sample buffer and boiled for 5 min. Protein extracts were resolved on SDS-PAGE (10% acrylamide) and transferred to a nitrocellulose Hybond ECL membrane (GE Healthcare, Pittsburgh, PA, USA) for probing with high-affinity rat monoclonal antibodies anti-HA (Roche, Basel, Switzerland) diluted 1:500, mouse anti-VSG221 diluted 1:500 and mouse anti-*Tb*SUMO diluted 1:500. Mouse monoclonal anti α-tubulin clone B-5-1-2 (Sigma-Aldrich, St. Louis, MO, USA) and anti-PABP [49] were used as loading controls. Horseradish peroxidase-conjugated goat anti-rat secondary antibody (Sigma-Aldrich) diluted 1:2000 was detected with SuperSignal® West Pico Chemiluminescent Substrate (Pierce, Rockford, IL, USA). Alexa Fluor® 790 AffiniPure goat anti-mouse IgG (H+L) or Alexa Fluor 680 AffiniPure goat anti-rabbit IgG (H+L) secondary antibodies (Jackson Immunoresearch Laboratories, West Grove, PA, USA) diluted 1:25000 were detected using an Odyssey laser-scanning system and quantified with Image Studio software (LI-COR Biosciences, Lincoln, NE, USA). Antibody signals were analyzed as integrated intensities of regions defined around the blots of interest.

### Growth curves

Parasite growth was evaluated by counting cell numbers daily by quadruplicate in a Neubauer haemocytometer. For doubling time calculation, parasites were maintained on exponential growth by diluting cultures every day to a density of 1×10^5^ cells/ml.

### Indirect immunofluorescence

BF cells were collected by centrifugation (1000 x g for 10 min), washed with TDB supplemented with glucose 20 mM and fixed with 4% paraformaldehyde (PFA) in PBS for 1 h. Parasites were allowed to bind to poly-L-lysine coated glass coverslips for 30 min and then incubated with 25 mM NH4Cl for 15 min. Permeabilization and blocking were performed with 3% bovine serum albumin (BSA), 0.5% saponin and 5% normal goat serum in PBS for 1 h. Mouse anti-*Tb*SUMO (1:500) [12], mouse anti-VSG221 (1:200), rabbit anti-VSG221 (1:500), mouse anti-EP FITC (1:500) (Cederlane Laboratories, Burlington, Canada) or anti-PAD1 [50] were used as primary antibodies. After washing with PBS, coverslips were incubated for 1 h with secondary antibodies diluted 1:1000 in 1% BSA:PBS (polyclonal goat anti-rabbit Alexa Fluor® 568 or polyclonal goat anti-mouse Alexa Fluor® 488 (Jackson)). Finally, coverslips were extensively washed and mounted using FluorSave reagent (Merck, Darmstadt, Germany). Nucleus and kinetoplast were visualized with 4,6-diamidino-2-phenylindole (DAPI) (Life Technologies). Samples were analyzed with an Eclipse 80i microscope (Nikon, Shinagawa, Japan).

### Flow cytometry

Parasites were collected by centrifugation, washed in TDB-glucose 20 mM and fixed with 1% PFA for 30 min at 4°C. After washing with PBS, cells were incubated with mouse anti-VSG221 (1:200) or mouse anti-EP FITC (1:500) diluted in 1% BSA:PBS for 30 min with gentle stirring. Parasites were then washed with PBS and resuspended in secondary antibodies diluted in 1% BSA:PBS (polyclonal goat anti-mouse Alexa Fluor® 488 (Jackson) diluted 1:200). Finally, cells were extensively washed with PBS an analyzed in a BD LSRFFortessa X-20 Cell Analyzer (BD, USA). Data analysis was performed using the FlowJo software (FlowJo LLC, Ashland, OR USA).

### *In vitro* differentiation

To promote differentiation to PF, BF parasites were resuspended at a density of 2.5×10^6^ cells/ml in SDM-79 media (Life Technologies) with 6 mM cis-aconitate (Sigma-Aldrich) at 28**°**C. Samples were collected by centrifugation at different time points and analyzed by immunofluorescence.

For stumpy differentiation, pleomorphic BF parasites from a mid-logarithmic growth phase culture were grown in semisolid HMI-9 agarose plates at low density (1×10^5^-1×10^6^ cells per plate). After 4 days of incubation at 37°C and 5% CO2, the stumpy-enriched population was harvested from the agarose plates by washing with HMI-9 and collected by centrifugation [51].

### Mice infections

Female BALB/c mice were inoculated intraperitoneally with 5000 parasites obtained from axenic culture in exponential growth. Parasitemias were obtained by counting cell number in blood with 0,83 %w/v ammonium chloride by quadruplicate in a Neubauer chamber under the light microscope (×400). Animal experiments were approved by the Committee on the Ethics of Animal Experiments of the Universidad Nacional de San Martin (CICUAE-UNSAM #11/17) and were carried out according to the recommendations of the Guide for the Care and Use of Laboratory Animals of the National Institutes of Health and the guidelines laid down by the Committee for the Care and Use of Animals for Experimentation.

### RNA isolation and gene expression analysis by RT-qPCR

Total RNA was isolated from ∼5×10^7^ BF parasites obtained from an axenic culture in exponential growth or blood of infected mice with Trizol Reagent, according to manufactureŕs instructions (Life Technologies). RNA integrity was assessed by electrophoresis and genomic DNA was eliminated with RQ1 DNAse (Promega, Madison, WI, USA) and subsequent chloroform extraction and ethanol precipitation. RNA was quantified by spectrophotometric assay with NanoDrop® system. cDNA was obtained from 3 µg of total RNA using 200 U Superscript II reverse transcriptase (Life Technologies) and 200 ng of random primers in a 20 µl total volume. Reactions were incubated at 42°C for 50 minutes and then at 70°C for 15 minutes. Quantitative PCR assays were carried out in the 7500 Real Time PCR System from Applied Biosystems using SensiFAST SYBR Lo-ROX Kit (BioLine, London, UK) and primers previously described [7, 52].

### ELISA assays

Microplates containing 96 wells (Thermo Scientific ImmunoPlates, MaxiSorp, Waltham, MA, USA) were coated overnight at 4°C with 5 µg/well of VSG221 purified as described by Cross [53] in PBS pH 7.4. Plates were washed 2 times with TBS-Tween (50 mM Tris-HCl (pH 7.6), 150 mM NaCl, 0.05% (v/v) Tween) and then blocked with 200 μl/well of buffer 1 (5% (w/v) skimmed milk in TBS-Tween) at room temperature (RT) for 1 h. The plates were then incubated for 1 h at RT with sera from infected mice diluted 1:5 in buffer 1. After washing 4 times with TBS-Tween, 100 μl of secondary antibody diluted in buffer 1 (peroxidase-conjugated goat anti-mouse IgM antibodies (Sigma-Aldrich)) were added and incubated at RT for 1 h. After additional washings with TBS-Tween the reaction was developed with tetramethylbenzidine for 15 min (TMB, Sigma-Aldrich) and stopped with 0.2 M sulphuric acid. Finally, absorbance values were measured at 450 nm in a microplate absorbance reader (FilterMax F5 Multimode, Molecular Devices, Sunnyvale, CA, USA).

## Acknowledgements

This work was supported by the National Agency for Promotion of Scientific and Technological Research, from the Argentinian Ministry of Science and Technology (ANPCyT, MinCyT), grants PICT-2016-0465 to VEA and PICT-2017-0140 to PAI. The funders had no role in study design, data collection and analysis, decision to publish, or preparation of the manuscript. We are grateful to G Montagna (IIBIO-UNSAM, Argentina) for helpful discussion and critical reading of the manuscript. We thank K. Matthews (Edinburgh) for providing PAD1 antibodies.

## Supp

**Figure S1:**
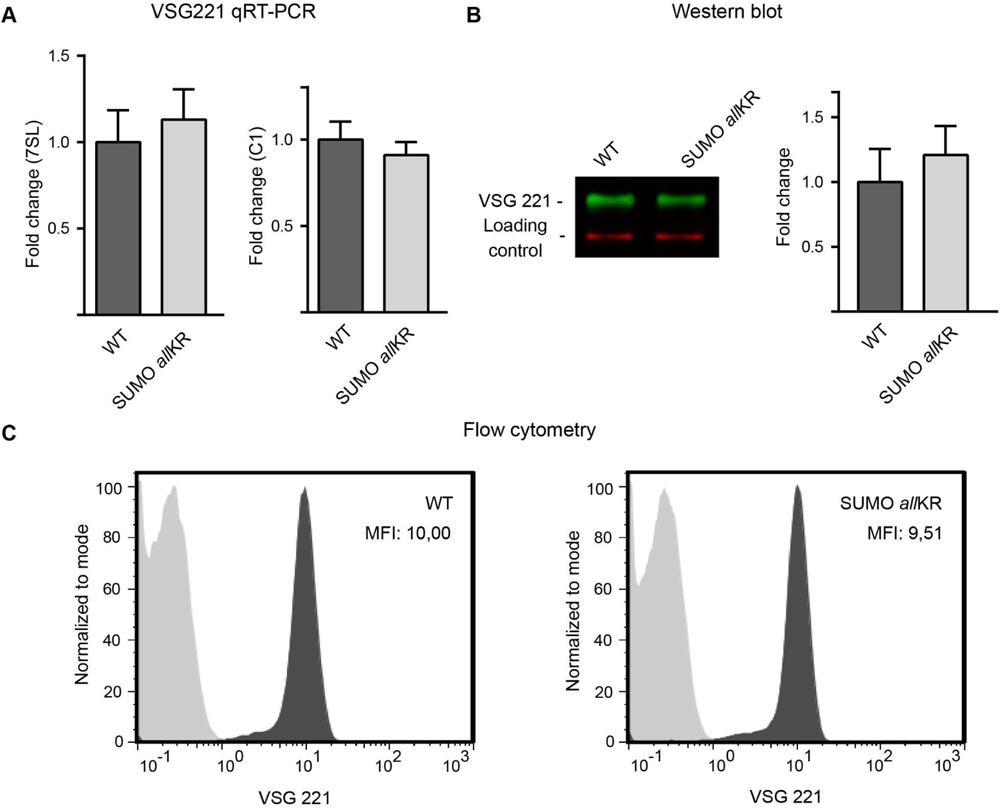
Analysis of VSG expression in SUMO chain competent *versus* SUMO chain mutant BF parasites. **(A)** Quantification of RNA transcript levels corresponding to the VSG221 gene expressed in wild type (WT) and SUMO *all*KR parasites. Transcript levels were determined using qRT-PCR and normalized against 7SL and C1. **(B)** VSG221 expression levels were evaluated by Western blot analysis in total cell extracts with anti-VSG221 antibodies and tubulin as a loading control. Experiments were performed at least in triplicates and one representative image is shown. Quantification of band intensities was performed using ImageJ. **(C)** The density of the VSG coat in WT and SUMO *all*KR parasites was analyzed in fixed cells with anti-VSG221 antibodies and flow cytometry. Approximately 50000 events were captured. Representative flow cytometry histograms normalized to mode are shown. MFI: median fluorescence intensity.

**Figure S2:**
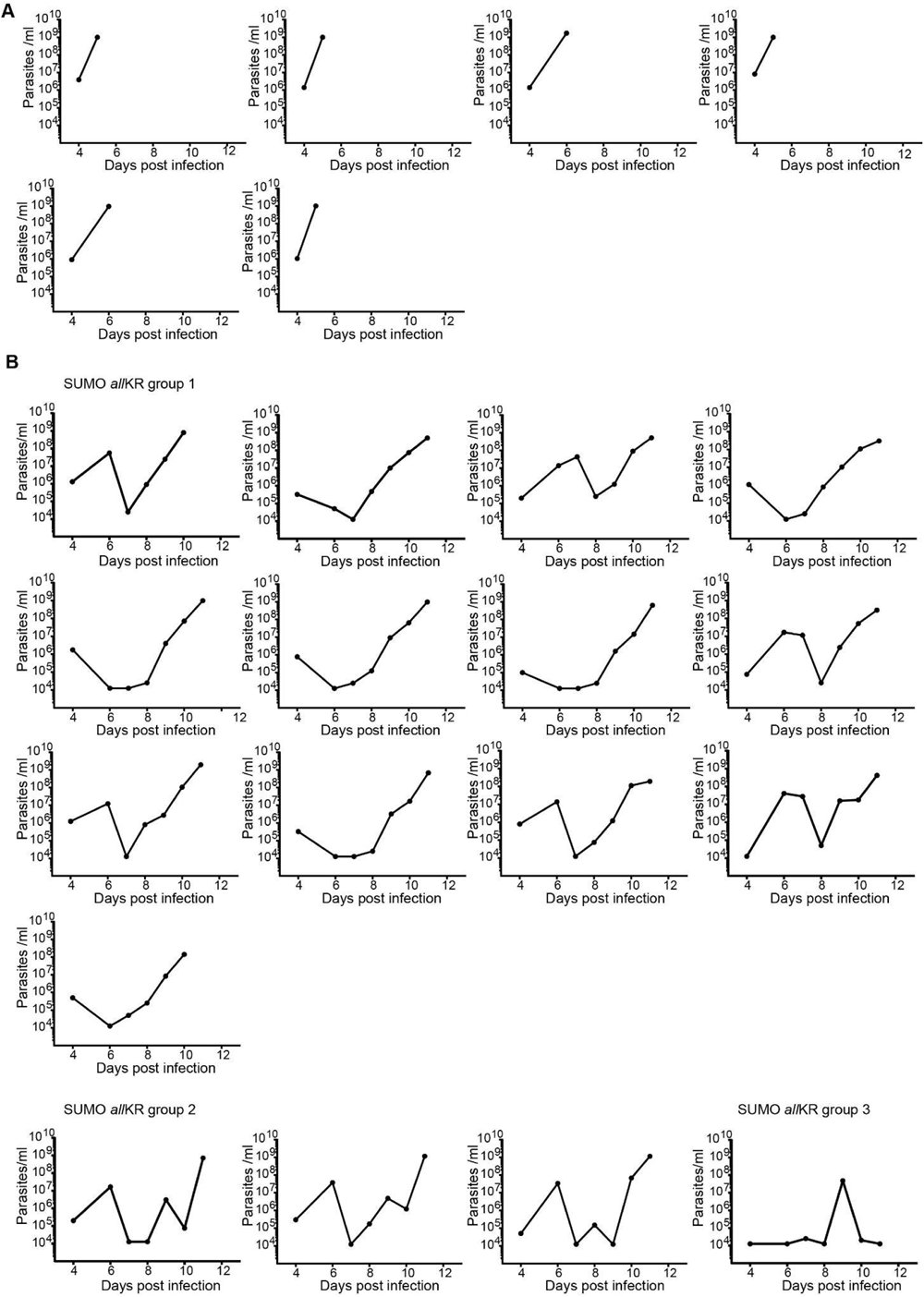
Mice infections with SUMO chain mutant BF parasites. Mice infected with SUMO *all*KR parasites display consecutives waves of parasitemia. Time course of parasitemia in mice infected with **(A)** wild type (WT) or **(B)** SUMO *all*KR parasites. Animals were grouped according to their different parasitemia profile. In the majority of the animals infected with SUMO chain mutants, parasites proliferated reaching the highest cell density approximately at 6 dpi (SUMO *all*KR group 1, n=13) after which the number of cells abruptly dropped at day 7 increasing again the day after. In other animals (SUMO *all*KR group 2, n=3) a pattern with two waves of parasitemia was observed and in one case (SUMO *all*KR group 3, n=1) the mouse cleared the infection and survived up to the end of the experiment. In general, all mice reached a maximum density of ∼5×10^7^-1×10^8^ parasites/ml before clearance, mostly in the second or third wave of parasitemia; while values around 5×10^8^-10^9^ parasites/ml led inexorably to death. In all cases a 3-log reduction in the number of parasites was observed, being the low parasitemia sustained for one or two consecutive days.

**Figure S3:**
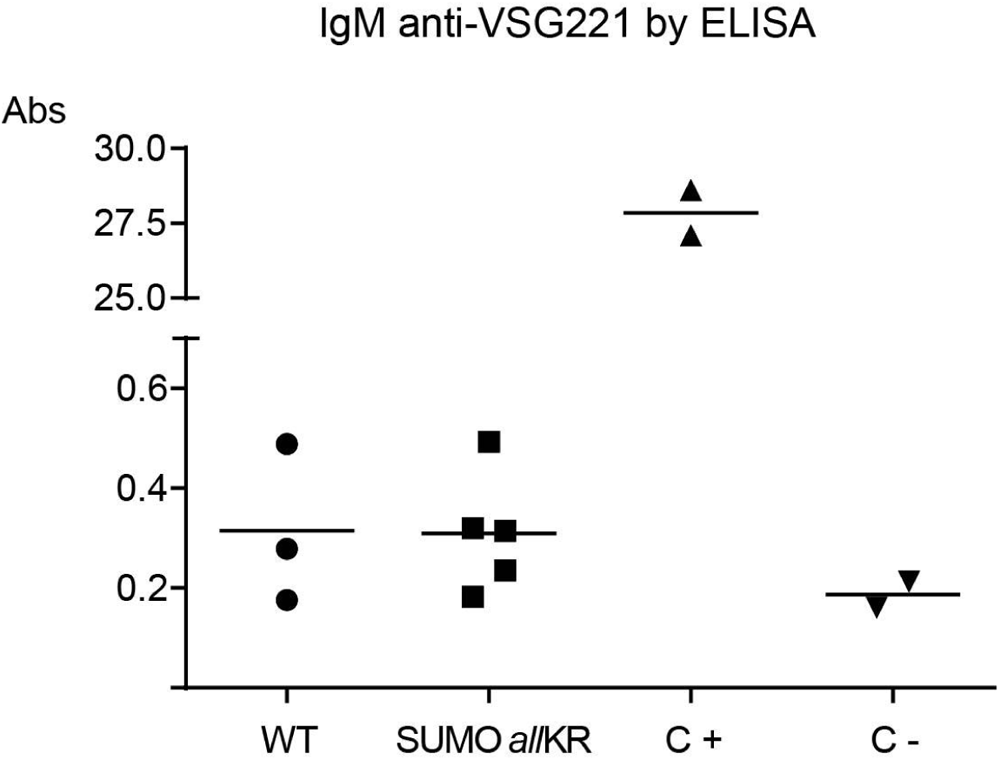
Host humoral immune response. Detection of IgM antibodies against VSG221 in the serum of mice infected with wild type (WT) or SUMO chain mutant (SUMO *all*KR) parasites at the first peak of parasitemia by ELISA. Serum samples were diluted 1:5, and absorbance values (Abs) were measured at 450 nm. C+: positive control; C-: negative control.

**Figure S4:**
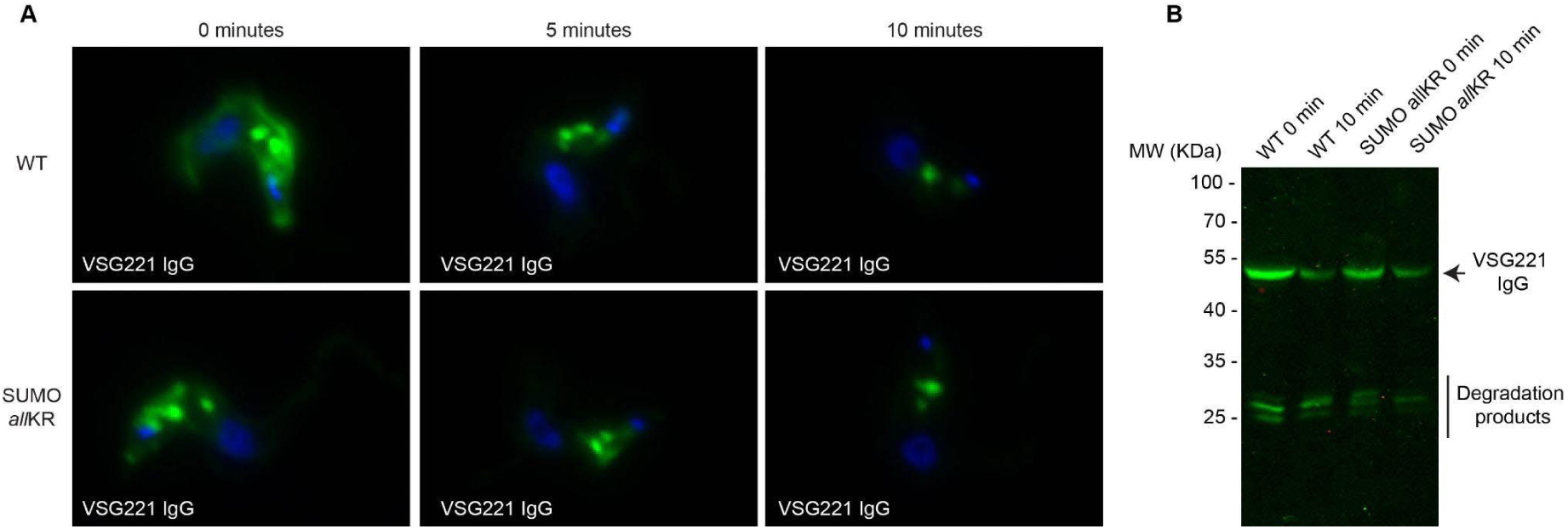
Degradation of anti-VSG IgG. WT or SUMO chain mutant (SUMO *all*KR) parasites were first incubated with anti-VSG221 IgG at 4°C in HMI-9, followed by an incubation at 37°C for 5 and 10 minutes. **(A)** Parasites were fixed and stained with anti-mouse-Alexa Fluor 488 (green) and DAPI (blue). **(B)** Whole-cell extracts (corresponding to 1×10^7^ cells) were obtained after the 10-minute incubation at 37°C and boiled in Laemmli’s sample buffer. Proteins were separated by SDS-PAGE and transferred to a nitrocellulose membrane, followed by immunoblotting with anti-mouse-Alexa Fluor 790.

**Figure S5:**
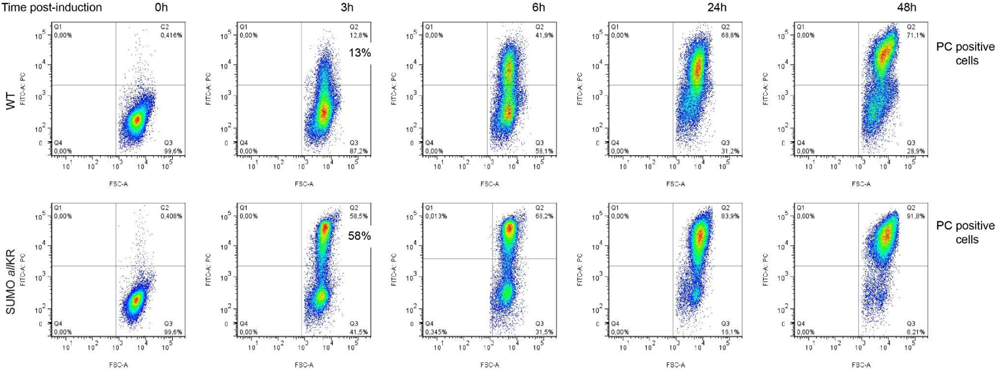
CA-induced differentiation of pleomorphic parasites. WT or SUMO *all*KR parasites were harvested from semisolid HMI-9 agarose plates and incubated with 6 mM cis-aconitate in SDM-79 at 28°C. Parasites were collected by centrifugation after different time points, fixed and stained with anti-EP FITC. Samples were analyzed by flow cytometry.

**Figure S6:**
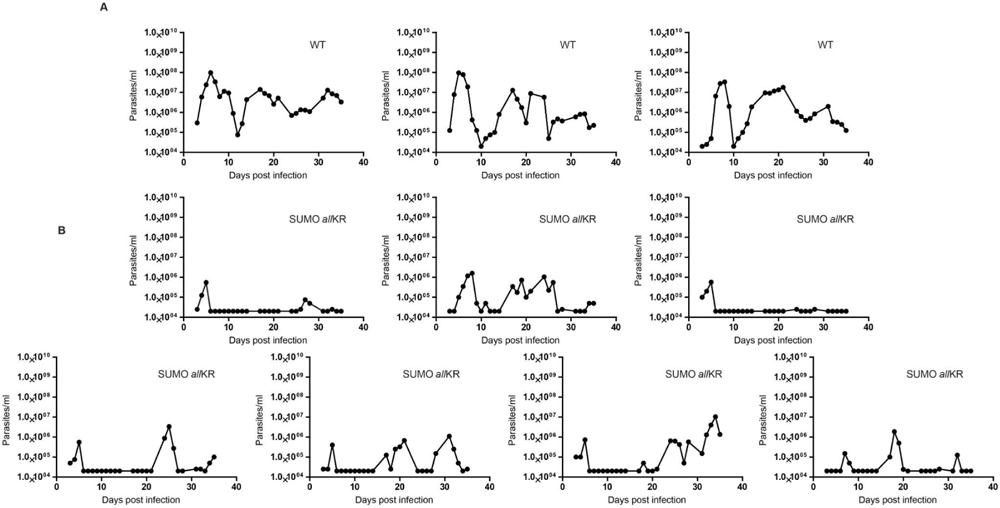
Mice infections with pleomorphic parasites. Time course parasitemia in mice infected with WT **(A)** or SUMO *all*KR **(B)** parasites.

